# mRNA induced expression of human angiotensin-converting enzyme 2 in mice for the study of the adaptive immune response to severe acute respiratory syndrome coronavirus 2

**DOI:** 10.1101/2020.08.07.241877

**Authors:** Mariah Hassert, Elizabeth Geerling, E. Taylor Stone, Tara L. Steffen, Madi S. Feldman, Jacob Class, Justin M. Richner, James D. Brien, Amelia K. Pinto

## Abstract

The novel human coronavirus, severe acute respiratory syndrome coronavirus 2 (SARS-CoV-2) has caused a pandemic resulting in nearly 20 million infections across the globe, as of August 2020. Critical to the rapid evaluation of vaccines and antivirals is the development of tractable animal models of infection. The use of common laboratory strains of mice to this end is hindered by significant divergence of the angiotensin-converting enzyme 2 (ACE2), which is the receptor required for entry of SARS-CoV-2. In the current study, we designed and utilized an mRNA-based transfection system to induce expression of the hACE2 receptor in order to confer entry of SARS-CoV-2 in otherwise non-permissive cells. By employing this expression system in an in vivo setting, we were able to interrogate the adaptive immune response to SARS-CoV-2 in type 1 interferon receptor deficient mice. In doing so, we showed that the T cell response to SARS-CoV-2 is enhanced when hACE2 is expressed during infection. Moreover, we demonstrated that these responses are preserved in memory and are boosted upon secondary infection. Interestingly, we did not observe an enhancement of SARS-CoV-2 specific antibody responses with hACE2 induction. Importantly, using this system, we functionally identified the CD4+ and CD8+ peptide epitopes targeted during SARS-CoV-2 infection in H2^b^ restricted mice. Antigen-specific CD8+ T cells in mice of this MHC haplotype primarily target peptides of the spike and membrane proteins, while the antigen-specific CD4+ T cells target peptides of the nucleocapsid, membrane, and spike proteins. The functional identification of these T cell epitopes will be critical for evaluation of vaccine efficacy in murine models of SARS-CoV-2. The use of this tractable expression system has the potential to be used in other instances of emerging infections in which the rapid development of an animal model is hindered by a lack of host susceptibility factors.

## Introduction

The pandemic level spread of severe acute respiratory syndrome coronavirus 2 (SARS-CoV-2), and the resulting outbreak of coronavirus disease 2019 (COVID-19) drives a need for the development of a range of novel animal models. SARS-CoV-2 results in a range of human disease phenotypes, most likely through different mechanisms[1]. This heterogeneity in human disease drives the need for a range of pre-clinical animal models necessary for the evaluation of direct acting therapeutics, host targeted therapeutics, the identification of vaccine correlates of protection, and a fundamental understanding of the drivers of pathology and disease. Each pre-clinical model will have its strengths and weaknesses.

The genome of the SARS-CoV-2 reference strain (Wuhan-Hu-1) (NC_045512.2) is ∼ 30 Kb and encodes four main structural genes: Spike, envelope (Env), membrane (Mem), nucleocapsid (NC) as well as 16 nonstructural proteins (nsp1-16) and multiple accessory proteins [2]. The reference strain of SARS CoV-2 shares approximately 82% identity with the SARS-CoV reference strain (NC_004718.3) at the nucleotide level, with greatest nucleotide differences occurring in the replicase, spike, and an accessory gene, 8a (GISAID, [3]). Interestingly, comparison at the amino acid level shows an approximate 77% identity, with the divergence of amino acids not localized to particular proteins shared between the two viruses. The similarities between SARS CoV and SARS CoV-2 have led to evidence of considerable serological cross-reactivity [4].

From the study of multiple human and animal CoVs, we know that the interaction between the spike glycoprotein and its cognate receptor is one of the main determinates regarding species and cellular tropism [5]. For HCoV-NL63, SARS-Co-V and SARS-CoV-2, the cognate receptor is the angiotensin-converting enzyme 2 (ACE2) [5-10]. ACE2 has been shown to be expressed in lungs, heart, kidneys and intestine and primarily functions as an enzyme controlling the maturation of angiotensin [11, 12]. Mice also express ACE2 and an amino acid alignment of the murine ACE2 (mACE2) with human ACE2 (hACE2) shows an approximate 81 percent identity between the two proteins (**Figure 1A**). As was shown for SARS-CoV [13], specific interactions between the spike glycoprotein, specifically the receptor binding motif of SARS-CoV-2 and hACE2, likely explain why SARS-CoV-2 infection of wild type mice does not occur [8]. Therefore, to establish a susceptible mouse model to study pathogenesis and immune responses to SARS-CoV-2, we must either alter the SARS-CoV-2 virus to recognize the mACE2[14] or express the hACE2 in mice [2, 15-21]. Various strategies have been employed by multiple groups to facilitate the expression of hACE2 in mice, from CRISPR/Cas9 mediated hACE2 transgenics [2, 15, 18-21], to adenovirus [16] or adeno associated virus (AAV) expressing hACE2 [17]. SARS-CoV-2 intranasal infections in these systems have achieved detectable virus in the lungs and trachea leading to viral pneumonia histologically [16, 17, 22]. Alternatively, targeted mutagenesis of the SARS-CoV-2 spike protein, facilitating enhanced interactions with murine ACE2, results in a similar phenotype [23]. Golden Syrian hamsters also offer a potential avenue to explore vaccines and therapeutics against SARS-CoV-2 and multiple groups have found that SARS-CoV-2 efficiently replicates in the nasal mucosa, bronchial epithelial cells and isolated areas of the lungs of hamsters and this viral replication results in pathological lesions similar to those found in COVID-19 patients [14, 24]. Interestingly, one of these groups has reported this as a transmission model [14]. However due to the lack of reagents at this time, hamster models have had limited utility in the studies of immune correlates of protection from infectious diseases [25].

**Figure 1:**
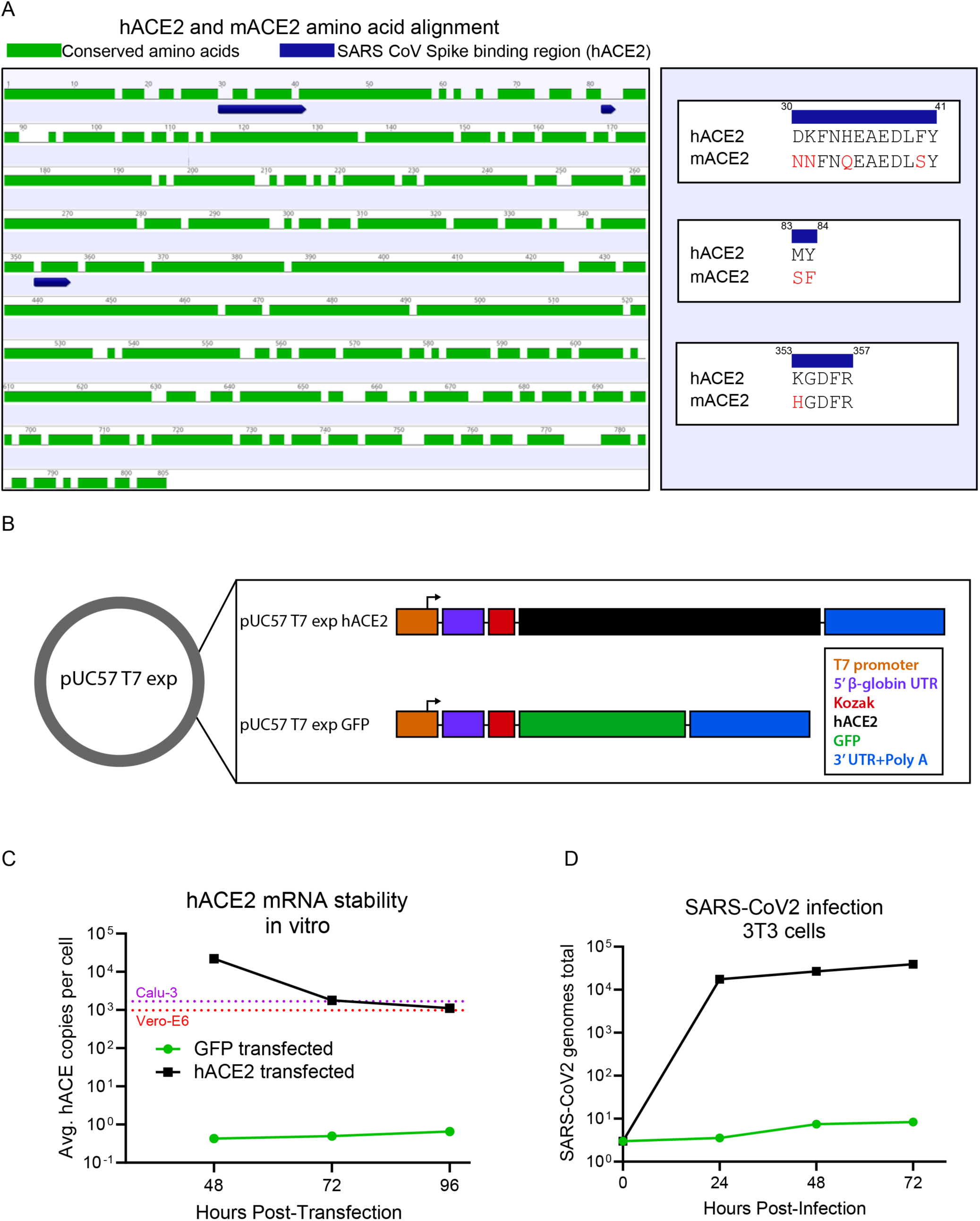
Transient expression of hACE2 in murine cells allows for SARS-CoV-2 entry. (**A**) Amino acid identity between hACE2 and mACE2. The amino acid sequences of human ACE2 (hACE2) (BAB40370) and murine ACE2 (mACE2) (NP_081562) were globally aligned using a BLOSUM62 cost matrix in the computational program Geneious. The two sequences showed 81.2% amino acid identity. Specific regions of interest included the amino acid residues important for SARS-CoV spike binding (and putative SARS-CoV-2 spike binding) (residues 30-41, 83-83, and 353-357). Multiple amino acid differences were noted in red in these critical regions between hACE2 and mACE2. (**B**) mRNA expression construct for induced expression of hACE2. T7 expression cassettes for hACE2 (and negative control GFP) were cloned into a pUC57 backbone by Gibson assembly. Each expression cassette includes a T7 promoter element, a 5’ β-globin UTR, Kozak sequence, CDS of each gene of interest (hACE2 or GFP), followed by a 3’ UTR and polyA tail. Following plasmid linearization and purification, mRNA was prepared in vitro using an ARCA T7 in vitro transcription kit. (**C**) In vitro stability of hACE2 mRNA. 2×10^6^ Murine 3T3 cells were transfected with 2 μg of either hACE2 or GFP mRNA and plated at a density of 5×10^6^ cells in each well of a 6 well dish. At 48, 72, and 96 hours post transfection, the stability of the hACE2 mRNA within the cells was assessed by qRT-PCR. (**D**) In vitro expression of hACE2 permits SARS-CoV-2 entry. 2×10^6^ Murine 3T3 cells were transfected with 2 μg of either hACE2 or GFP mRNA and plated at a density of 5×10^6^ cells in each well of a 6 well dish. 24 hours post transfection, cells were infected with SARS-CoV-2. At 24, 48, and 72 hours post infection, SARS-CoV-2 RNA was quantified from the 3T3 cell pellets by qRT-PCR.

By developing an mRNA delivery platform to express hACE2 in mice, we have established a unique and highly adaptable murine model for studying SARS-CoV-2 infection. Using this model, we were able to characterize the murine adaptive immune response generated against SARS-CoV-2. We noted many significant benefits of the mRNA system for the development of a susceptible animal model for SARS-CoV-2, including rapid development of a susceptible mouse. The transfected RNA is highly durable, as we were able to detect mRNA expression in culture at high levels for at least 4 days following transfection. The transfection treatment can also be given repeatedly as the mRNA is poorly immunogenic. Utilizing the C57BL/6 type 1 interferon receptor deficient (Ifnar1^-/-^) mice transfected with hACE2 mRNA, we have demonstrated that these mice are susceptible to a transient infection and can generate SARS-CoV-2 specific adaptive immune responses. We have been able use this system to identify both neutralizing antibodies and virus specific T cell responses. In this study we have identified nine CD8^+^ and six CD4^+^ H2^b^ restricted SARS CoV-2-specific T cell epitopes. These results demonstrate the development of a novel animal model for SARS-CoV-2 where hACE2 is expressed by mRNA and its utility in studying the adaptive immune responses to SARS CoV-2.

## Results and Discussion

To generate a construct which could produce robust expression of hACE2 mRNA, we used a pUC57 vector backbone consisting of four core components: 1) a type II T7 promoter for a high transcription rate *in vitro* and the production of RNA transcripts with homogeneous 5’ and 3’ termini; 2) a beta globin 5’UTR for optimized mRNA expression within mammalian cells; 3) 3’ UTR from alpha globin for mRNA stability and 4) A 153 base adenosine nucleotide stretch as the polyA tail. Into this backbone, we cloned either the coding sequence for hACE2 or green fluorescent protein (GFP), as a control (**Figure 1B**). mRNA from either construct was then generated using a T7 ARCA in vitro transcription reaction. To demonstrate the stability of transcribed hACE2 mRNA, murine fibroblast 3T3 cells were transfected with the in vitro transcribed and purified GFP or hACE2 mRNA. At 48, 72 and 96 hours post-transfection, expression of hACE2 was measured by qRT-PCR (**Figure 1C**). For hACE2, the qRT-PCR data revealed that we were able to detect stable transfected hACE2 mRNA in the 3T3 cells for four days, with only a minimal decline in expression levels over that time period. For at least 96 hours post-transfection, the expression level of hACE2 in this hACE2 transfected cell line was similar to the mRNA expression of hACE2 we detected highly SARS-CoV-2 susceptible Calu-3 and Vero-E6 cell lines **(Figure 1C**). With the cells transfected with the GFP expressing mRNA construct, we performed flow cytometry and demonstrated that we could achieve greater than 90 percent transfection efficiency, and that the GFP expression was maintain for at least 72 hours (**Supplemental Figure 1**). We attempted to measure hACE2 expression on hACE2 mRNA transfected 3T3 cells by flow cytometry in a similar manner. However, due to high antibody cross-reactivity between hACE2 and the murine cognate mACE2, after evaluation of six commercially available monoclonal and polyclonal antibodies, we concluded that mACE2 and hACE2 could not be distinguished in this manner and were unable to confirm hACE2 expression in our transfected 3T3 cells (data not shown). These results demonstrated that we were able to generate an mRNA vector expression system and deliver hACE2 to non-hACE2 expressing murine 3T3 cells, resulting in RNA expression levels similar to that observed in susceptible cell lines (Calu-3 and Vero-E6), and that the expression was stable for at least 96 hours.

We next confirmed that expression of hACE2 by our mRNA construct conferred susceptibility of SARS-CoV-2 in murine cells by infecting the hACE2 or GFP transfected 3T3 cells with SARS-CoV-2 (**Figure 1D)**. The hACE2 or GFP mRNA transfected cells were each plated into separate 6 well plates, and 24 hours after transfection, they were infected with SARS-CoV-2 at a multiplicity of infection (MOI) of 0.01. The infected cells were harvested at 24, 48, and 72 hours post infection and the viral genomes were quantified by qRT-PCR. The 3T3 cells transfected with the hACE2 had more virus detected at every time point post infection with approximately four logs higher viral genome copies within the cells as compared to the cells that had been transfected with the GFP expressing mRNA (**Figure 1D**). These results demonstrate that the transfection of murine cells with hACE2 mRNA confers susceptibility to infection with SARS-CoV-2.

### Adaptive immune response to SARS-CoV-2

Understanding the role of the antigen specific T cell and antibody response against SARS-CoV-2 is critical for the development of safe and effective vaccines. Studies with murine hepatitis virus (MHV) A59, a natural mouse coronavirus pathogen [26], concluded that a T cell response in combination with an antibody response was critical for controlling a coronavirus infection [27]. Additionally, studies into SARS-CoV and MERS-CoV suggest that the receptor binding domain (RBD) of the spike glycoprotein is likely the most effective target for current antibody-based vaccines against SARS-CoV-2. However, these SARS-CoV and MERS-CoV vaccine studies also noted that there is a concern that a vaccine directed solely against the spike glycoprotein will induce viral escape [28, 29], and a successful vaccine should contain both a strong T cell response for early viral control, and neutralizing antibody for viral clearance [30-32]. Therefore, understanding the role of the T cell as well as the antibody response against SARS CoV-2 will be critical for the development, testing and evaluation of future vaccines.

To begin to identify the immune correlates of protection we used the hACE2 mRNA transfection system to study the adaptive immune response to SARS-CoV-2 in type I interferon receptor 1 deficient (Ifnar1^-/-^) mice (**Figure 2**). We chose to use the Ifnar1^-/-^ mice, because, like other viral infections, coronaviruses encode genes which dampen or block the type I IFN response creating a more susceptible environment for the establishment of infection [33, 34]. Studies with multiple viruses have shown that the loss of IFNAR, and therefore the significant dampening of IFN stimulating gene production, results in IFNAR deficient cells being highly susceptible to viral infections [35-44]. Therefore, for the reported SARS-CoV-2 animal studies, we have used Ifnar1^-/-^ mice.

**Figure 2:**
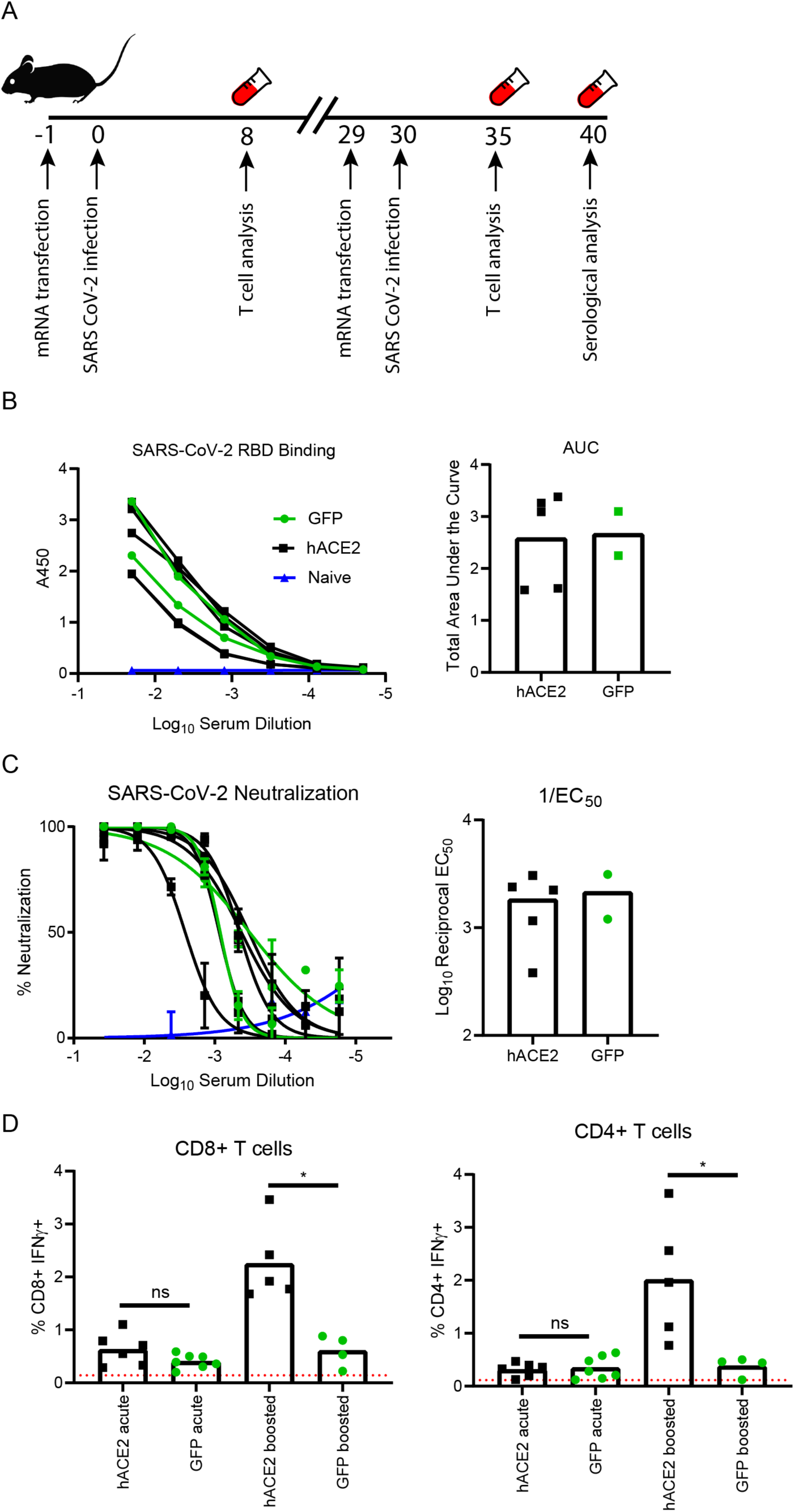
In vivo transfection of hACE2 mRNA yields enhanced CD4^+^ and CD8^+^ T cell responses. (**A**) Experimental design. 8-week-old Ifnar1-/- mice were transfected in vivo with 10 μg of either GFP or hACE2 mRNA. 24 hours post transfection, mice were infected with SARS-CoV-2. At day 8 post infection, blood was collected for acute phase T cell stimulation assays. At days 29 post initial infection, the mice were again transfected in vivo with 10 μg of either GFP or hACE2 mRNA and infected 24 hours later. 5 days post boost, blood was collected for memory recall T cell stimulation assays and serum was collected for serological analysis. (**B**) Spike receptor binding domain ELISA. Recombinant SARS-CoV-2 spike RBD protein was used to coat an immunosorbent plate. Serum from transfected and infected mice was serially diluted and used to determine RBD binding potential by absorbance at 450nm with increasing serial dilution and area under the curve calculation. (**C**) Neutralization potential of polyclonal sera. Serum from transfected and SARS-CoV-2 infected mice was serially diluted and incubated with ∼100 focus forming units of SARS-CoV-2 to allow complexes to form. Virus-serum complexes were then overlayed on a Vero-WHO monolayer and allowed to infect for 24 hours, at which point the plates were fixed and developed (see materials and methods). Neutralization was determined by enumerating a reduction in infectious particles with increased serum concentration and determining the EC_50_. (**D**) Global T cell responses during SARS-CoV-2 infection. 8 days post initial transfection with hACE2 mRNA or GFP mRNA and infection and 5 days post boost, blood lymphocytes were collected from each mouse and stimulated with anti-CD3 for 6 hours in the presence of brefeldin A. Following stimulation, cells were washed and stained with anti CD19, CD4, CD8, IFNγ and TNFα. The frequency of responding CD8+ and CD4+ T cells was demonstrated by quantifying the frequency of CD8+ or CD4+ T cells producing IFNγ. Statistical significance was determined by Mann-Whitney test (p=0.0159).

In these studies, we followed a prime boost strategy (**Figure 2A**), similar to what would be done for a vaccine, so to enhance the detection of the immune response to SARS-CoV-2 in the Ifnar1^-/-^ mice. First, the mice were transfected with 10 μg of mRNA encoding either hACE2 or GFP as a control. Twenty-four hours after mRNA in vivo delivery, we infected the hACE2 or GFP transfected Ifnar1^-/-^ mice with 5×10^4^ focus forming units (FFU) per mouse of SARS-CoV-2 administered both intravenously (IV) and intranasally (IN). Blood was collected eight days post infection for acute phase T cell analysis. After day 21 post primary infection, the Ifnar1^-/-^ mice received a second hACE2 or GFP mRNA transfection, followed 24 hours later by a boost with 5×10^4^ FFU per mouse of SARS-CoV-2, administered both IV and IN route. Five days post boost, blood was collected for T cell and antibody analysis.

To examine the antibody response directed against SARS-CoV-2, we performed an indirect ELISA against the receptor binding domain (RBD) of the SARS-CoV-2 spike protein, a target of the neutralizing antibody response in both mice and humans [16, 45-47]. To quantify the anti-RBD antibody response, we compared the polyclonal sera from naïve Ifnar1^-/-^ mice, to that of the SARS-CoV-2 infected Ifnar1^-/-^ mice transfected with either the hACE2 or GFP mRNA (**Figure 2B**). We noted that only the Ifnar1^-/-^ mice that had been infected with SARS CoV-2 were able to bind RBD. As expected, we saw no RBD binding from sera collected from naïve mice. However, analysis of the area under the curve (AUC) showed there were no detectible differences between the SARS CoV-2 infected Ifnar1^-/-^ mice that had received GFP and hACE2 mice at either time point.

While there were SARS-CoV-2 RBD specific antibodies present in both the infected GFP and hACE2 transfected mice, we were interested to determine if there was a difference in the neutralization capacity of the antibody response generated in the hACE2 transfected mice as compared to the GFP transfected controls. To test this, we performed focus reduction neutralization (FRNT) assays on the serum samples (**Figure 2C**). Briefly, serum from each mouse was serially diluted and combined with ∼100 FFU of SARS-CoV-2 for one hour before the antibody-virus complex was added to a monolayer of Vero cells. After 24 hours of infection, the cells were fixed and stained with anti-SARS polyclonal guinea pig sera then stained with horseradish peroxidase-conjugated goat anti-guinea pig IgG. Foci from infected cells were visualized and counted using an EliSpot plate reader. Similar to the results of the ELISA, serum from naïve Ifnar1^-/-^ mice was not able to neutralize SARS-CoV-2 (**Figure 2C**). Analysis of the neutralization curves and FRNT_50_ values from the serum collected showed that both the hACE2 and GFP transfected Ifnar1^-/-^ mice were able to generate neutralizing antibodies, and there were no differences in the neutralization abilities of the serum collected at this time point(average EC_50_ values of 0.00096 and 0.00032, respectively). From the RBD ELISA data and FRNT data, we concluded that Ifnar1^-/-^ mice infected with SARS CoV-2 can generate an antibody response to the virus and this response is independent of the hACE2 expression.

Using the prime boost protocol outline in **Figure 2A** we examined the CD8^+^ T cell and CD4^+^ T cells from the blood after primary infection, and boost, by intracellular cytokine staining for the detection of interferon gamma (IFN-γ) in response to anti-CD3 stimulation (**Figure 2D and Supplemental Figure 2**). During the acute infection the IFN-γ production in both the CD4^+^ and CD8^+^ T cells isolated from both the GFP and hACE2 transfected group were only slightly above the background of unstimulated cells isolated from the same group of mice and were not significantly different from one another. However, following the second infection with SARS-CoV-2 the IFN-γ response was significantly higher in the Ifnar1^-/-^mice that were transfected with hACE2 (average CD8^+^ response= 2.25% and average CD4+ response= 2.01%) as compared to GFP (average CD8^+^ response= 0.61% and average CD4+ response= 0.38%) (p=0.0159). Importantly, we did not see a significant increase in the IFN-γ responses from the CD4^+^ or CD8^+^ T cell in the boosted GFP mice as compared to the GFP mice following primary infection. This suggests that GFP transfected mice did not have an amnestic T cell response upon boost with SARS-CoV-2.

The results of these studies indicate that the expression of hACE2 on the murine cells allows SARS-CoV-2 to enter the murine cells and undergo viral replication. This viral replication provides the antigen processing and presentation pathways access to the SARS-CoV-2 proteins. The proteins can then be either directly or cross-presented to T cells for the development of a robust response. This result is supported both by *in vitro* studies in the 3T3 cells (**Figure 1D**) and the *in vivo* T cell and FRNT studies (**Figure 2**). Combining these studies suggests that hACE2 expression delivered by the mRNA construct allows virus entry and replication which is required for the production of viral epitopes needed to stimulate a T cell response.

### Viral replication

To test whether mRNA expression of hACE2 would allow the detection of infectious virus output, we followed a similar protocol to the one we had used in the adaptive immune studies where we administered 10 μg of the hACE2 mRNA or control GFP mRNA to Ifnar1^-/-^ mice one day prior to infection. 24 hours post transfection, the Ifnar1^-/-^ mice receiving either the hACE2 or the GFP mRNA constructs were infected with 5×10^4^ FFU per mouse using a combined IV and IN route. Three days post SARS-CoV-2 infection, the mice were harvested and viral titers were measured in the brain, kidney, spleen, liver, lung and whole blood both by qRT-PCR and by focus forming assay (FFA). We were able to detect viral genome copies in the transfected Ifnar1^-/-^ mice, however we did not detect differences in viral genome copies between the hACE2 and the GFP recipient mice (**Supplemental Figure 3A-F**). Additionally, we did not detect infectious virus in any organ from either group of mice by focus forming assay (FFA) (data not shown).

**Figure 3:**
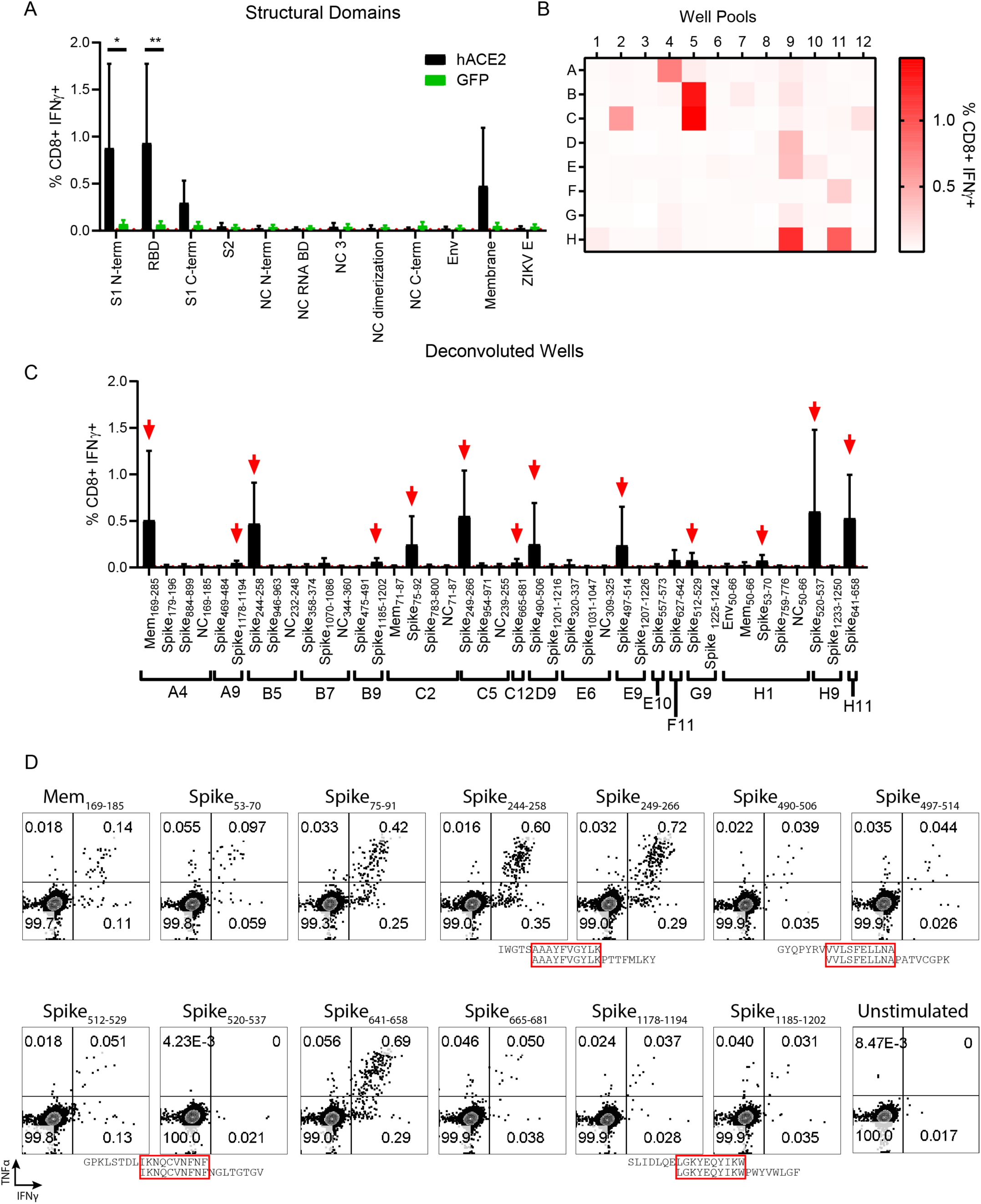
In vivo transfection of hACE2 mRNA permits the detection and functional mapping of SARS-CoV-2 specific CD8+ T cell responses. (**A**) CD8+ T cell responses to pooled peptide domains. Each peptide library was demarcated into peptides contained in functional domains of each protein and peptides contained in each domain were pooled into equimolar pools (11 total pools). 5 days post infection with SARS-CoV-2 following transfection with either hACE2 or GFP mRNA, splenocytes were harvested and stimulated for 6 hours with each peptide pool in the presence of brefeldin A. After stimulation, cells were stained for flow cytometry to evaluate the frequency of responsive CD8+ T cells by IFNγ expression. (**B**) CD8+ T cell responses to smaller well peptide pools. Each library was incorporated into multiple 96-well plate formats (**Supplemental Figure 4**). Within the same layout, wells from the plates were pooled such that all A1 peptides were pooled, all A2 peptides, etc. maintaining the 96-well plate format, but reducing the overall number of samples that needed to be screened. 5 days post boosted infection following transfection with hACE2 mRNA, splenocytes were harvested and stimulated with each peptide pool in the presence of brefeldin A. The frequency of IFNγ+ CD8+ T cells— the magnitude of which represents responsiveness to a peptide in the pool, is enumerated in a heat map format as the average responses of 3 mice. (**C**) Once potential well hits were identified, the peptides contained in each well were deconvoluted and used individually to stimulate splenocytes from hACE2 transfected, SARS-CoV-2 infected mice. 13 potential epitopes were identified (marked with red arrows) as defined by the frequency of IFNγ+ CD8+ T cells being at least 2-fold above background (stimulated with vehicle) in at least 3 of the 4 mice screened. (**D**) Representative flow cytometry plots displaying IFNγ and TNFα expression in CD8+ T cells for each putative epitope in comparison to a vehicle control. Due to the overlapping nature of the peptide library, Spike_244-258_ and Spike_249-266_, likely are demonstrating responsiveness to the same peptide epitope. 3 other instances of this phenomenon are denoted with red boxes surrounding the amino acid sequence overlap. Statistical significance was determined by Mann-Whitney test (*p=.0159, **p=0.0079)

While these studies suggest that the hACE2 mRNA transfection into the Ifnar1^-/-^ mice did not increase the susceptibility of the Ifnar1^-/-^ mice to SARS-CoV-2 it is possible that the dose and route of infection as well as the harvest time points we chose for these studies were not optimal for detection of infectious virus. Previous studies have shown that administration of higher doses of virus by the intranasal route did lead to the detection of virus in mice which had been induced to express hACE2 through the administration of an adeno-associated virus vector or recombinant Adenovirus [2, 15-22]. While in the current study we chose low dose administration of virus to focus on immune responses future studies with higher doses of virus, similar to those published in other mouse models [2, 15-22] can be administered to determine the utility of this model for SARS-CoV-2 pathogenesis. Additionally, future studies with alternate routes will be useful in determining if the delivery of mRNA expressing hACE2 can be used to develop a mouse model to study virus pathogenesis in specific tissues.

### SARS-CoV-2 CD8^+^ and CD4^+^T cell epitope identification

While the mRNA delivery of hACE2 followed by infection with 5×10^4^ FFU of SARS-CoV-2 did not allow our group to study viral pathogenesis in Ifnar1^-/-^ mice, the delivery of mRNA hACE2 to our murine model allowed for the induction of a strong CD4^+^ and CD8^+^ T cell response to SARS-CoV-2 (**Figure 2D**). With this result, we chose to use our SARS-CoV-2 murine model to identify the SARS-CoV-2-specific CD8^+^ and CD4^+^ T cell responses using a peptide library screening method (**Figures 3 and 4)**. We chose this approach, as it has worked previously for identifying Zika virus (ZIKV) T cell epitopes using a ZIKV peptide library in C57BL/6 mice [36, 48]. For SARS-CoV-2, we screened peptide libraries spanning the structural genes including the spike (BEI: NR-2669), nucleocapsid (NC) (BEI: NR-52404), envelope (Env) (BEI:NR-52405) and Membrane (Mem) (BEI: NR-52403). Each of the peptide arrays of 12-mer to 20-mers overlapped by approximately 10 amino acids and spanned the entire length of each protein. For the initial peptide screening, a SARS-CoV Urbani strain (GenBank: AY278741) Spike peptide library (BEI: NR-2669) was used in place of SARS-CoV-2 Spike (BEI: NR-52402) due to lack of reagent availability. For the Env, NC, and Mem screening assays, peptide libraries generated from of sequences from SARS-CoV-2 USA-WA1/2020 strain were used. Each peptide from the library was reconstituted and added to an individual well in a 96 well plate with each gene plated into a separate 96 well plate (**Supplemental Figure 4 and Supplemental Table 1**). The resulting peptide library was spread across five plates with a total of 269 coronavirus structural peptides, that were screened in the initial studies.

**Table 1:** Functional identification of SARS-CoV-2 CD8+ T cell epitopes. H2-b restricted mice were transfected with hACE2 mRNA and infected with SARS-CoV-2, rested for 30 days and boosted with SARS-CoV-2. At 5 days post boost, splenocytes from 4 mice were harvested and stimulated with peptide or vehicle control in the presence of brefeldin A for 6 hours. Cells were stained for flow cytometry using anti-CD19, CD4, CD8, and IFNγ. CD8+ T cells were defined as CD19 negative, CD8 positive, and CD4 negative. Antigen-specific cells were identified as CD8 T cells producing IFNγ. ^**1**^Peptide sequences are named based on the protein they are contained within, followed by the number of the first amino acid residue of the peptide in the context of the full protein, to the last amino acid residue. Sequences contained within the “spike” peptide library correspond with SARS-CoV spike due to library availability. ^**2**^Exact amino acid residues of peptide used to stimulate splenocytes. ^**3**^Name of the SARS-CoV-2 predicted peptide epitopes. ^**4**^9-mer or 8-mer predicted SARS-CoV-2 epitope predicted based on location and identity of larger peptide sequences used to stimulate splenocytes and of anchor residues in SARS-CoV-2. ^**4**^Predicted MHC haplotype based on conserved anchor residues. ^**5**^Fold over background frequency of IFNγ+ CD8 T cells. Background is defined as the frequency of IFNγ+ CD8 T cells in a well stimulated with vehicle control.

| Library Peptide Name<sup>1</sup> | Library Peptide Sequence<sup>2</sup> | SARS-CoV-2 Predicted Peptide Name<sup>3</sup> | SARS-CoV-2 Predicted Epitope<sup>4</sup> | Predicted MHC Haplotype<sup>5</sup> | Fold Over Background<sup>6</sup> |  |  |  |
| --- | --- | --- | --- | --- | --- | --- | --- | --- |
|  |  |  |  |  | Mouse 1 | Mouse 2 | Mouse 3 | Mouse 4 |
| Mem<sub>169-185</sub> | TVATSRTLSTYYKLGASQ | Mem<sub>174-181</sub> | RTLSTYYKL | H2-K<sup>b</sup> | 8.33 | 81.00 | 13.85 | 26.45 |
| Spike<sub>53-70</sub> | YLTQDLFLPFYSNVTGFH | Spike<sub>54-62</sub> | LFLPFSSNV | H2-K<sup>b</sup> | 5.20 | 3.90 | 10.08 | 4.07 |
| Spike<sub>75-91</sub> | TFGNPVIPIFKDGIYFAA | Spike<sub>76-983</sub> | TKRFDNPV | H2-D<sup>b</sup> | 22.33 | 13.20 | 6.84 | 5.01 |
| Spike<sub>244-258</sub> | IWGTSAAAYFVGYLK | Spike<sub>263-270</sub> | AAYYVGYL | H2-K<sup>b</sup> | 31.67 | 37.00 | 21.91 | 26.20 |
| Spike<sub>249-266</sub> | AAAYFVGYLKPTTFMLKY | Spike<sub>263-270</sub> | AAYYVGYL | H2-K<sup>b</sup> | 33.67 | 47.00 | 26.20 | 37.28 |
| Spike<sub>490-506</sub> | GYQPYRVVLSFELLNA | Spike<sub>511-518</sub> | VVLSFELL | H2-K<sup>b</sup> | 2.47 | 45.70 | ND | 2.22 |
| Spike<sub>497-514</sub> | VVLSFELLNAPATVCGPK | Spike<sub>511-518</sub> | VVLSFELL | H2-K<sup>b</sup> | 2.33 | 43.00 | ND | 4.00 |
| Spike<sub>512-529</sub> | GPKLSTDLIKNQCVNFNF | Spike<sub>539-546</sub> | VNFNFNGL | H2-K<sup>b</sup> | 6.03 | 5.05 | 2.28 | 1.07 |
| Spike<sub>520-537</sub> | IKNQCVNFNFNGLTGTGV | Spike<sub>539-546</sub> | VNFNFNGL | H2-K<sup>b</sup> | 0.70 | 95.50 | 68.01 | 50.38 |
| Spike<sub>641-658</sub> | HVDTSYECDIPIGAGICA | Spike<sub>658-665</sub> | NSYECDIP | H2-K<sup>b</sup> | 32.67 | 44.00 | 46.85 | 17.13 |
| Spike<sub>665-681</sub> | LLRSTSQKSIVAYTMSL | Spike<sub>691-699</sub> | SIIAYTMSL | H2-K<sup>b</sup> | 2.93 | 4.20 | 4.53 | 3.12 |
| Spike<sub>1178-1194</sub> | SLIDLQELGKYEQYIKW | Spike<sub>1203-1211</sub> | LGKYEQYIK | H2-K<sup>b</sup> | 2.17 | 3.70 | 4.69 | 6.55 |
| Spike<sub>1185-1202</sub> | LGKYEQYIKWPWYVWLGF | Spike<sub>1203-1211</sub> | LGKYEQYIK | H2-K<sup>b</sup> | 2.20 | 5.55 | 12.32 | 3.18 |

**Figure 4:**
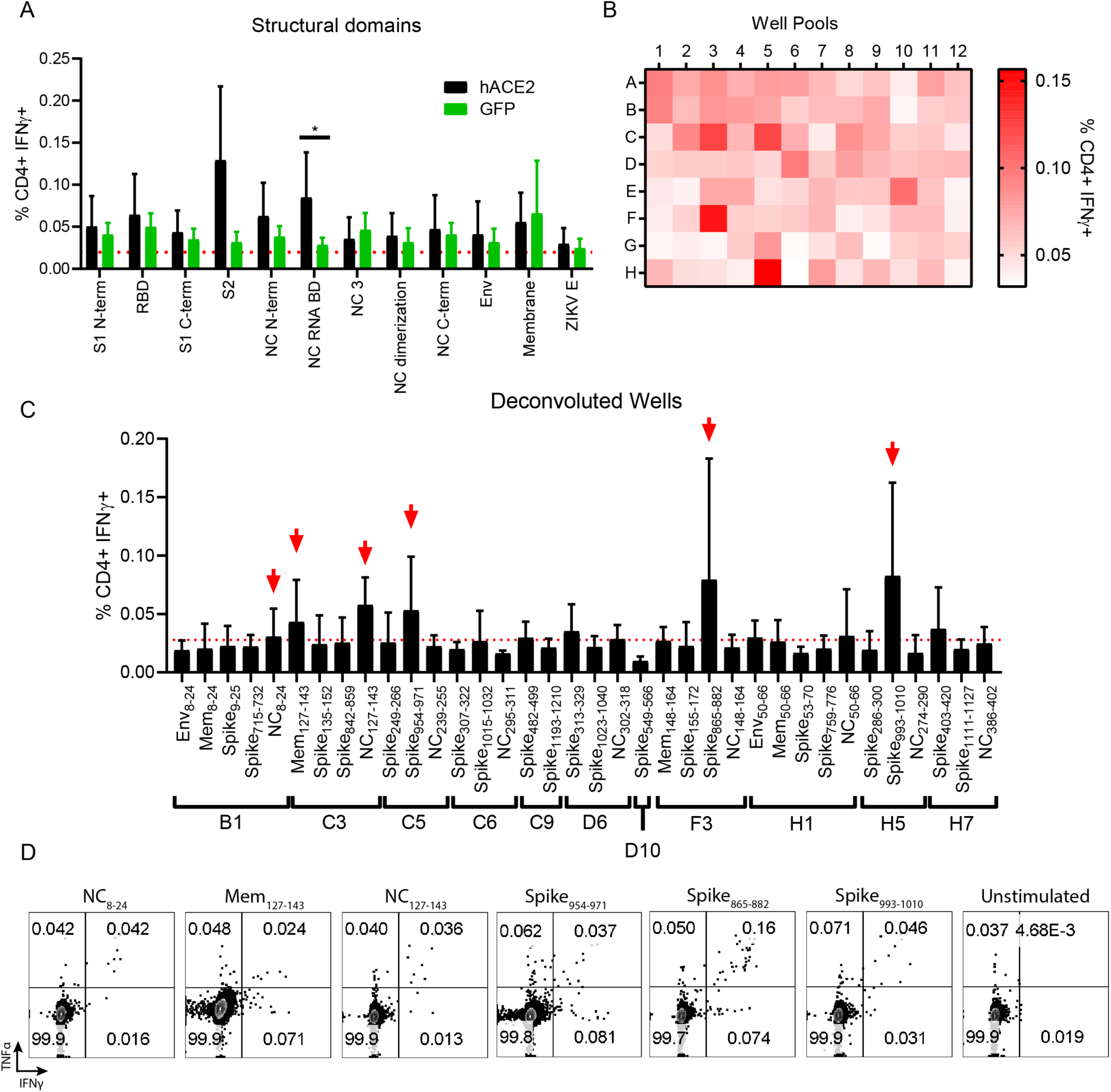
In vivo transfection of hACE2 mRNA permits the detection and functional mapping of SARS-CoV-2 specific CD4+ T cell responses. (**A**) CD4+ T cell responses to pooled peptide domains. 5 days post infection with SARS-CoV-2 following transfection with either hACE2 or GFP mRNA, splenocytes were harvested and stimulated for 6 hours with each domain peptide pool in the presence of brefeldin A (as in Figure 3A). After stimulation, cells were stained for flow cytometry to evaluate the frequency of responsive CD4+ T cells by IFNγ expression. (**B**) CD4+ T cell responses to smaller “well” peptide pools. As was done in Figure 3B, each library was incorporated into multiple 96-well plate formats and pooled. 5 days post boost following transfection with hACE2 mRNA, splenocytes were harvested and stimulated with each peptide pool in the presence of brefeldin A. The frequency of IFNγ+ CD4+ T cells is enumerated in a heat map format as the average responses of 3 mice. (**C**) Once potential “well” hits were identified, the peptides contained in each well were deconvoluted and used individually to stimulate splenocytes from hACE2 transfected, SARS-CoV-2 infected mice. 6 potential epitopes were identified (marked with red arrows) as defined by the frequency of IFNγ+ CD4+ T cells being at least 2-fold above background (stimulated with vehicle) in at least 3 of the 4 mice screened. (**D**) Representative flow cytometry plots displaying IFNγ and TNFα expression in CD4+ T cells for each putative epitope in comparison to a vehicle control. Statistical significance was determined by Mann-Whitney test (*p=.0317).

To identify the epitope targets of the SARS-CoV-2 specific CD8^+^ T cells in our primary screen, we followed a similar prime-boost infection strategy detailed in **Figure 2A** where we delivered hACE2 or GFP mRNA followed by infection with 5×10^4^ FFU of SARS-CoV-2. After 21 days, we again transfected mice with hACE2 mRNA followed 24 hours later with 5×10^4^ FFU of SARS-CoV-2. At day 5 post SARS-CoV-2 boost, mice were sacrificed and spleens were homogenized into single cell suspensions. The single cell suspensions of splenocytes were plated into 96 well plates and stimulated for six hours in the presence of Brefeldin A (BFA) and either a gene specific peptide pool (**Figure 3A**). After the stimulation, splenocytes were stained with the cell surface antibodies, α-CD8, α-CD4, and α-CD19, then stained intracellularly with antibodies against the mouse cytokines interferon gamma (IFN-γ) and tumor necrosis factor-α (TNF-α) as shown in **Supplemental Figure 2**. To identify antigen experienced CD8^+^ T cells from the SARS-CoV-2 boosted animals, we compared the results of the peptide stimulation against unstimulated cells and a ZIKV envelope peptide pool as a negative control, and anti-CD3 (clone 45-2C11) stimulated cells as a positive control. Pools that demonstrated an IFN-γ response that was more than two-fold over background in the majority of mice screened were brought forward to identify possible SAR-CoV-2 epitopes. Using this approach, it became clear that the majority of the CD8+ T cell response in these H2^b^ restricted mice targets peptides within the spike and membrane proteins (**Figure 3A**). As demonstrated in **Figure 2D**, transfection with hACE2 mRNA prior to infection and boost enhanced SARS-CoV-2 specific T cell response. Therefore, for specific epitope identification, hACE2 transfected animals were used moving forward.

To further narrow down potential epitope targets, peptides were pooled in an alternative manner in which pools of one to five peptides were generated by combining the same well from each of the five plates from the aliquoted library shown in **Supplemental Figure 4 and Supplemental Table 1** and were similarly used to stimulate splenocytes of hACE2 mRNA transfected and SARS-CoV-2 boosted mice (**Figure 3B**). Using this approach 17 well pools screened positive for potential CD8^+^ T cell epitopes present (**Figure 3B**). Using this information, the peptide pools were deconvoluted and individual multi-mer peptides from each presumptive positive well was evaluated (**Figure 3C**). The individual peptides within the peptide pools were screened by repeating the hACE2 transfection and SARS-CoV-2 boosting strategy in the Ifnar1^-/-^ mice (**Figure 2A**). With this approach, we identified 13 wells that contained peptides which induced a cytokine response that was greater than 2-fold above background in the majority of mice screened (**Table 1**).

Using the information from the peptide screen we predicted potential optimal SARS-CoV-2 peptide sequences for each of the epitopes based on known K^b^ and D^b^ anchor residues. Due to the overlapping nature of the screening libraries used, we hypothesize that 8 of the library peptide hits are actually 4 epitopes which are present in multiple wells (**Figure 3D**). To demonstrate that the responses to each peptide were polyfunctional, we assessed IFN-γ and TNF-α expression. All nine of the epitopes we identified were polyfunctional based upon co-expression of both IFN-γ and TNF-α in response to stimulation. We named the SARS-CoV-2 peptide epitopes with the abbreviated name of the viral protein followed by the number of the amino acid based upon the SARS-CoV-2 open reading frame for each gene, for example Mem_169-185_ would mean the epitope began at the 169^th^ amino acid in the Mem open reading frame (**Table 1**).

The identical approach, using the same mice, was concurrently used to characterize the CD4^+^ T cell responses to SARS-CoV-2 (**Figure 4**). Stimulation of the CD4^+^ T cells with the gene specific peptide pool (**Figure 4A**) and the pool peptides from the combined five plates (**Figure 4B)** showed weak but detectible responses spread across all 12 protein domains evaluated. While the signal to noise ratio of the CD4^+^ T cell response was lower than what we had observed for the CD8^+^ T cells, the responses from the protein domains pooled peptides (**Figure 4A**) suggested that the strongest CD4^+^ T cell response was located in the S2 region of the spike protein, with responses also seen in the RBD and NC pools. Similar to the CD8^+^ T cell responses, we saw an enhancement of the SARS-CoV-2 specific CD4^+^ T cell responses as a result of hACE2 mRNA transfection. By evaluating the smaller well pools of peptides consisting of 1-5 pooled peptides, we identified 11 positive wells, where the IFN-γ responses to the peptide pool was greater than 2-fold over background in multiple animals (**Figure 4B**). By deconvoluting those wells to individual peptides, we identified six novel SARS-CoV-2 reactive CD4^+^ T cell epitopes in C57BL/6 mice **(Figure 4C and Table 2)**. Similar to what we observed for the peptide stimulated CD8^+^ T cells, the CD4^+^ T cells stimulated with the individual peptide epitopes were able to make both IFN-γ and TNF-α in response to peptide stimulation **(Figure 4D)**.

**Table 2:** Functional identification of SARS-CoV-2 CD4+ T cell epitopes. H2-b restricted mice were infected with SARS-CoV-2, rested for 30 days and boosted with SARS-CoV-2. At 5 days post boost, splenocytes from 4 mice were harvested and stimulated with peptide or vehicle control in the presence of brefeldin A for 6 hours. Cells were stained for flow cytometry using anti-CD19, CD4, CD8, and IFNγ. CD4+ T cells were defined as CD19 negative, CD8 negative, and CD4 positive. Antigen-specific cells were identified as CD4 T cells producing IFNγ. ^**1**^Peptide sequences are named based on the protein they are contained within, followed by the number of the first amino acid residue of the peptide in the context of the full protein to the last amino acid residue. Sequences contained within the “Spike” peptide library correspond with SARS-CoV Spike due to library availability. ^**2**^Exact amino acid residues of peptide used to stimulate splenocytes. ^**3**^Analogous SARS-CoV-2 peptide name in instances where SARS-CoV peptide library had to be used due to reagent availability. ^**4**^Analogous SARS-CoV-2 peptide sequence in instances where SARS-CoV peptide library had to be used due to reagent availability. ^**5**^Predicted MHC haplotype based on conserved anchor residues. ^**6**^Fold over background frequency of IFNγ+ CD8 T cells. Background is defined as the frequency of IFNγ+ CD4 T cells in a well stimulated with vehicle control.

| Library Peptide<sup>1</sup> | Library Peptide Sequence<sup>2</sup> | SARS-CoV-2 Predicted Peptide<sup>3</sup> | SARS-CoV-2 Analogous Sequence<sup>4</sup> | MHC<sup>5</sup> | Fold Over Background<sup>6</sup> |  |  |  |
| --- | --- | --- | --- | --- | --- | --- | --- | --- |
|  |  |  |  |  | Mouse 1 | Mouse 2 | Mouse 3 | Mouse 4 |
| Mem<sub>127-143</sub> | TILTRPLLESELVIGAV | Mem<sub>127-143</sub> | TILTRPLLESELVIGAV | I-A<sup>b</sup> | 2.32 | 1.09 | 2.71 | 4.69 |
| NC<sub>127-143</sub> | KDGIWVATEGALNTPK | NC<sub>127-143</sub> | KDGIWVATEGALNTPK | I-A<sup>b</sup> | 1.76 | 2.05 | 5.79 | 10.36 |
| NC<sub>8-24</sub> | NQRNAPRITFGGPSDST | NC<sub>8-24</sub> | NQRNAPRITFGGPSDST | I-A<sup>b</sup> | ND | 2.43 | 0.97 | 7.30 |
| Spike<sub>865-882</sub> | TAGWTFGAGAALQIPFAM | Spike<sub>883-900</sub> | TSGWTFGAGAALQIPFAM | I-A<sup>b</sup> | 0.73 | 9.81 | 2.86 | 4.77 |
| Spike<sub>954-971</sub> | AISSVLNDILSRDKVEA | Spike<sub>972-989</sub> | AISSVLNDILSRDKVEA | I-A<sup>b</sup> | 2.88 | 2.22 | 1.73 | 6.09 |
| Spike<sub>993-1010</sub> | QLIRAAEIRASANLAATK | Spike<sub>1011-1028</sub> | QLIRAAEIRASANLAATK | I-A<sup>b</sup> | 4.80 | 3.23 | 2.28 | 8.80 |

There is a significant gap in our knowledge concerning infection and correlates of protection for coronaviruses, including the novel coronavirus SARS-CoV-2. Much of that deficit is due to the lack of tractable small animal models in which the reagents necessary to study T and B cell responses are available. Our goal was to use an mRNA delivery system to express hACE2, the putative receptor for SARS CoV-2 in mice, to generate a small animal model to study the adaptive immune response to SARS CoV-2 infection. By developing a highly tractable hACE2 expression system, we have generated the potential for an animal model which can be rapidly used by multiple groups to study aspects of disease, transmission, and immune protection for SARS-CoV-2. Importantly, this system is adaptable and has the potential to be employed for use in the rapid development of animal models of emerging infectious diseases which lack necessary host susceptibility factors. In the present study, we utilized this system to understand the adaptive immune response during SARS-CoV-2 infection in Ifnar^1-/-^ mice. We were able to demonstrate that multiple administrations of hACE2-encoding mRNA can be used to detect enhanced CD4^+^ and CD8^+^ T cell responses to SARS-CoV-2 during a prime and boost infection (**Figure 2**).

Additionally, by using this approach in conjunction with an overlapping peptide library to stimulate these T cells, we identified nine SARS-CoV-2 CD8^+^ T cell epitopes and six CD4^+^ T cell epitopes which are H2^b^ restricted. In doing so, we expanded the knowledge within the field of the adaptive immune response to SARS-CoV-2, moving much-needed pre-clinical animal models for SARS-CoV-2 and COVID-19 forward.

## Materials and Methods

### Ethics statement-

All animal studies were conducted in accordance with the Guide for Care and Use of Laboratory Animals of the National Institutes of Health and approved by the Saint Louis University Animal Care and Use Committee (IACUC; protocol 2771).

### Generation of hACE2 and GFP constructs-

GFP and hACE2 expression constructs were cloned by Gibson assembly into a pUC57 vector (sequences available). The plasmids were linearized downstream of the 3’ UTR and polyA tail by XbaI digestion. The linearized plasmids were purified and utilized as templates in a T7 ARCA in vitro transcription reaction (New England Biolabs). The mRNA product was then purified using an Invitrogen Purelink RNA mini kit according to the manufacturer’s instructions. Transcript length and quality was confirmed by RNA bleach gel.

### Virus and cells-

SARS-CoV-2 (Isolate USA-WA1/2020) was obtained from BEI (catalog NR-52281) A p1 stock was grown in African green monkey kidney cells (Vero-E6) purchased from ATCC using this initial seed stock. A p2 stock was then grown from this p1 stock by infecting Vero-E6 cells at an MOI of 0.01 in complete DMEM and harvested at 96 hours post infection. In vitro transfection and infection experiments were completed using murine fibroblast 3T3 cells cultured in complete DMEM and were obtained from American Type Culture Collection (ATCC CRL-1658).

### In vitro transfection-

3T3 cells were transfected with either hACE2 or GFP mRNA using an Amaxa cell line nucleofector L kit (Lonza; catalog number VCA-1005) according to the manufacturer’s instructions. 5×10^5^ transfected cells were plated in each well of a 6-well dish. hACE2 mRNA stability was measured by qRT-PCR at 48, 72, and 96 hours post transfection. Stability of GFP expression was confirmed by flow cytometry at 24, 48, and 72 hours post transfection. Twenty-four hours post transfection, susceptibility of the cells was evaluated by infecting with SARS-CoV-2 at an MOI of 0.01 in 0.5 ml of DMEM. At 24, 48, and 72 hours post infection, media supernatant was removed from the wells and RNA was extracted from the cells using an Invitrogen Purelink RNA mini kit according to the manufacturer’s instructions.

### Mice, in vivo transfection, and infections-

Type 1 interferon receptor deficient mice (Ifnar1-/-) were purchased from Jackson laboratories and maintained as a colony in a pathogen-free mouse facility at Saint Louis University-School of Medicine. Eight-week-old Ifnar1-/- mice were transfected with 10 μg of RNA using Polyplus in vivo-jet RNA in vivo transfection reagent prepared according to the manufacturer’s instructions and administered via intravenous (IV) and intranasal (IN) combination route (100 μl and 20 μl, respectively). 24 hours following transfections, mice were infected with 5×10^4^ focus forming units (FFU) of SARS-CoV-2 via IV and IN combination route (100 μl and 20 μl, respectively). A subset of mice (n=6 GFP and n=7 hACE2) were used to quantify viral burden at 3 days post infection. Mice were administered a lethal dose of ketamine/xylazine cocktail and perfused with 20 ml of PBS. Blood was collected into RNAsol BD, and viral RNA was extracted according to the manufacturer’s instructions. Spleen, liver, kidney, brain, and lung tissues were collected into Sarstedt tubes and snap frozen. Organs were homogenized in DMEM using a bead beater and viral RNA was extracted from the organ lysates using TriReagent RT. For antibody and T cell experiments, mice were transfected with 10 μg of RNA using Polyplus in vivo-jet RNA in vivo transfection reagent prepared according to the manufacturer’s instructions and administered via intravenous (IV) and intranasal (IN) combination route. 24 hours following transfections, mice were infected with 5×10^4^ focus forming units (FFU) of SARS-CoV-2 via IV and IN combination route. At day 8 post infection, blood was collected for acute phase T cell analysis. 30 days following initial infection, mice were again transfected with RNA and infected in the same manner. Blood was collected at 5 days post boost for anti-CD3 T cell stimulation experiments and again at day 10 post boost for serological analysis. Splenocytes from mice were harvested at 5 days post boost for T cell epitope mapping.

### qRT-PCR-

hACE2 expression was measured by qRT-PCR using Taqman primer and probe sets from IDT (assay ID Hs.PT.58.27645939). SARS-CoV-2 viral burden was measured by qRT-PCR using Taqman primer and probe sets from IDT with the following sequences: Forward 5’ GAC CCC AAA ATC AGC GAA AT 3’, Reverse 5’ TCT GGT TAC TGC CAG TTG AAT CTG 3’, Probe 5’ ACC CCG CAT TAC GTT TGG TGG ACC 3’. Synthesized hACE2 RNA was used as a copy control to quantify the number of hACE2 molecules present in each sample. Similarly, a SARS-CoV-2 copy number control (available from BEI) was used to quantify SARS-CoV-2 genomes.

### T cell stimulation-

For anti-CD3 stimulation of peripheral blood lymphocytes, blood was collected via submandibular cheek bleed directly into alkaline lysis buffer. After red blood cell lysis, cells were washed twice with complete RPMI media (10% FBS, 1X HEPES, and 1X beta-mercaptoethanol) and resuspended in complete RPMI. Cells were then split between 2 wells of a 96-well round bottom plate and stimulated for 6 hours at 5% CO_2_ and 37°C in the presence of 10 μg/ml brefeldin A with 5 μg/ml of anti-CD3 (clone 2C11) or water or a pool of Zika virus envelope peptides as negative controls. For peptide stimulation of splenocytes, spleens were harvested into complete RPMI medium from mice 5 days post SARS-CoV-2 boost. Spleens were ground over a 70μm filter and then washed with 10ml of RPMI. Approximately 5×10^5^ cells were plated per well in a 96-well round bottom plate and stimulated for 6 hours at 5% CO_2_ and 37°C in the presence of 10 μg/ml brefeldin A and 50 μg/ml of each peptide or peptide pools.

### Flow cytometry-

Following stimulation of lymphocytes, cells were washed once with PBS and stained overnight in PBS at 4°C for the following surface antigens: CD4 (clone RM-4-5), CD8α (clone 53-6.7), and CD19 (clone 1D3). Cells were washed in PBS, then fixed in 2% paraformaldehyde at 4°C for 10 minutes. After fixation, cells were permeabilized with 0.5% saponin and stained in 0.5% saponin at 4°C for 1 hour for the following intracellular antigens: TNFα (clone Mab11) and IFNγ (clone B27). After intracellular staining, cells were washed with 0.5% saponin followed by PBS. The cells were analyzed by flow cytometry using an Attune NxT focusing flow cytometer. For analysis, CD4+ T cells were gated on lymphocytes, CD19 negative, CD4 positive and CD8 negative cells. CD8+ T cells were gated on lymphocytes, CD19 negative, CD4 negative and CD8 positive. Antigen specific cells were then identified as producing IFNγ and/or TNFα at greater than 2-fold over cells stimulated with just a vehicle control.

### SARS-CoV-2 Receptor binding domain ELISA-

To determine the binding potential of polyclonal sera from SARS-CoV-2 infected mice to the hACE2 receptor binding domain (RBD) of SARS-CoV-2, maxisorp ELISA plates were coated overnight at 4°C with 1 μg/ml of recombinant SARS-CoV-2 RBD protein in carbonate buffer. The following day, the plates were blocked with PBS, 5% BSA, and 0.5% Tween for 2 hours at room temperature prior to being washed. Serum from each mouse was serially diluted and added to each well and allowed to incubate for 1 hour at room temperature prior to being washed. Horseradish peroxidase conjugated goat-anti-human IgG secondary antibody was added and allowed to incubate for 1 hour at room temperature prior to being washed. TMB enhanced substrate was added and allowed to incubate in the dark at room temperature for 15 minutes prior to quenching with 1N HCl. Following quenching, absorbance of the plate was read at 450 nanometers using a BioTek Epoch plate reader.

### Focus reduction neutralization assay-

Serum from each mouse was serially diluted in DMEM containing 5% FBS and combined with ∼100 focus forming units (FFU) of SARS-CoV-2 and allowed to complex at 37°C and 5% CO_2_ for 1 hour in a 96-well round bottom plate. The antibody-virus complex was then added to each well of a 96-well flat bottom plate containing a monolayer of Vero WHO cells. Following 1 hour of incubation at 37°C and 5% CO_2_, the cells were overlayed with 2% methylcellulose and returned to the incubator. After 24 hours of infection, the cells were fixed with 5% electron microscopy grade paraformaldehyde in PBS for 15 minutes at room temperature. The cells adherent to the plate were then rinsed with PBS and permeabilized with 0.05% Triton-X in PBS. Foci of infected Vero cells were stained with anti-SARS polyclonal guinea pig sera (BEI) overnight at 4°C and washed 3 times with 0.05% Triton-X in PBS. Cells were then stained with horseradish peroxidase conjugated goat anti-guinea pig IgG for 2 hours a room temperature. Cells were washed again with 0.05% Triton-X in PBS prior to the addition of TrueBlue KPL peroxidase substrate, which allows the visualization of infected foci as blue spots. The foci were visualized and counted using an ImmunoSpot CTL Elispot plate reader.

### Peptide library-

SARS-CoV-2 and SARS structural protein peptide libraries were obtained from BEI Resources. A SARS spike (catalog NR-2669) peptide library was used in place of SARS-CoV-2 spike due to limited reagent availability. The library consisted of a 169-peptide array of 15-mers to 20-mers overlapping by approximately 10 amino acids and spanning the length of the SARS Urbani strain S protein (GenBank: AY278741). The SARS CoV-2 envelope peptide library (catalog NR-52405) consisted of an array of 10 peptides ranging from 12-mer to 15-mers and overlapping by 10 amino acids spanning the length of the envelope protein of SARS-CoV-2 USA-WA1/2020 strain (GenPept: QHO60596). The SARS-CoV-2 membrane peptide library (catalog NR-52403) consisted of an array of 31 12-mer to 17-mer peptides overlapping by 10 amino acids spanning the length of the membrane protein of SARS-CoV-2 USA-WA1/2020 strain (GenPept: QHO60597). The SARS-CoV-2 nucleocapsid peptide library (catalog NR-52404) consisted of an array of 59 peptides ranging from 13-mers to 17-mers with 10 amino acids of overlap spanning the length of the nucleocapsid protein of SARS-CoV-2 USA-WA1/2020 strain (GenPept: QHO60601). Amino acid sequence information can be found in **Supplemental Table 1**. Each peptide came in a lyophilized vial and was reconstituted in 90% DMSO to 10 mg/ml and oriented in a 96 well plate format. During reconstitution, no peptides were noted as insoluble. After reconstitution, subsets of peptides were consolidated to form 11 peptide pools containing various regions or predicted subdomains of each protein (N-terminal region of S1, receptor binding domain, C-terminal region of S1, S2, N-terminal region of nucleocapsid, RNA binding domain of nucleocapsid, nucleocapsid group 3, dimerization domain of nucleocapsid, C-terminal region of nucleocapsid, envelope, and membrane) (**Supplemental Figure 4**). In addition, smaller peptide pools were formed by combining analogously oriented wells from each peptide plate (e.g. all A1 peptides are pooled) (**Supplemental Figure 4**).

## Acknowledgements

*Financial support-*This work was supported by Saint Louis University COVID-19 research Seed Funding to awarded to AKP and awarded to JDB, National institutes of Health grant F31 AI152460-01 from the National Institute of Allergy and Infectious Diseases (NIAID) awarded to MH.

## Author contributions

MH, JDB, and AKP conceptualized the work and wrote and edited the manuscript. Expression construct design and cloning was completed by MH. In vitro transcription and RNA purification was completed by MH, JC, and JR. Viral stocks were grown and quantified by MH. In vitro transfection and SARS-CoV-2 infection experiments were completed by MH. In vivo transfection and SARS-CoV-2 infections were completed by MH and LG. Viral burden in vivo was determined by LG. SARS-CoV-2 RBD ELISAs were completed by TLS. Focus Reduction Neutralization Tests were completed by ETS. Evaluation of the T cell response to SARS-CoV-2 was completed by MH. All authors reviewed and approved the final version of the manuscript.

## Figure Legends

**Supplemental Figure 1: Protein expression stability from expression constructs**. 2×10^6^ Murine 3T3 cells were transfected with 2 μg of either hACE2 or GFP mRNA and plated at a density of 5×10^6^ cells in each well of a 6 well dish. At 24, 48, and 72 hours GFP expression was evaluated by flow cytometry compared to untransfected cells.

**Supplemental Figure 2: T cell epitope mapping gating strategy**. T cells were defined by a lymphocyte gate based on size and granularity and CD19 negative. T cells were further classified as CD8+ or CD4+ T cells by staining CD8+/CD4- or CD4+/CD8-respectively. Antigen responsive T cells were defined by IFN-γ expression.

**Supplemental Figure 3: Viral burden in Ifnar^1-/-^ mice at day 3 post infection** Ifnar^1-/-^ mice were transfected with 10 μg of either GFP or hACE2 RNA. 24 hours following transfections, mice were infected with 5×10^4^ focus forming units (FFU) of SARS-CoV-2 via IV and IN combination route (100 μl and 20 μl, respectively). n=6 GFP and n=7 hACE2 were used to quantify viral burden at 3 days post infection in the lungs (**A**), spleen (**B**), liver (**C**), kidney (**D**), brain (**E**), and whole blood (**F**) by qRT-PCR.

**Supplemental Figure 4: Plate maps of SARS-CoV and SARS-CoV-2 peptide libraries**. Peptide libraries spanning the SARS-CoV or SARS-CoV-2 structural proteins were obtained from BEI (**Supplemental Table 1**). Every 12-17-mer peptide came in a lyophilized vial and was reconstituted in 90% DMSO to 10 mg/ml and oriented in a 96 well plate format. Subsets of peptides were consolidated to form 11 peptide pools containing various regions or predicted subdomains of each protein (N-terminal region of S1, receptor binding domain, C-terminal region of S1, S2, N-terminal region of nucleocapsid, RNA binding domain of nucleocapsid, nucleocapsid group 3, dimerization domain of nucleocapsid, C-terminal region of nucleocapsid, envelope, and membrane). To aid in identification, peptide pools of 1-5 peptides were also made consisting of the identical well of each plate (e.g. all A1 wells were pooled).

