## Supplementary figures and images for "mRNA induced expression of human angiotensin-converting enzyme 2 in mice for the study of the adaptive immune response to severe acute respiratory syndrome coronavirus 2"

### Supplemental Figure 1

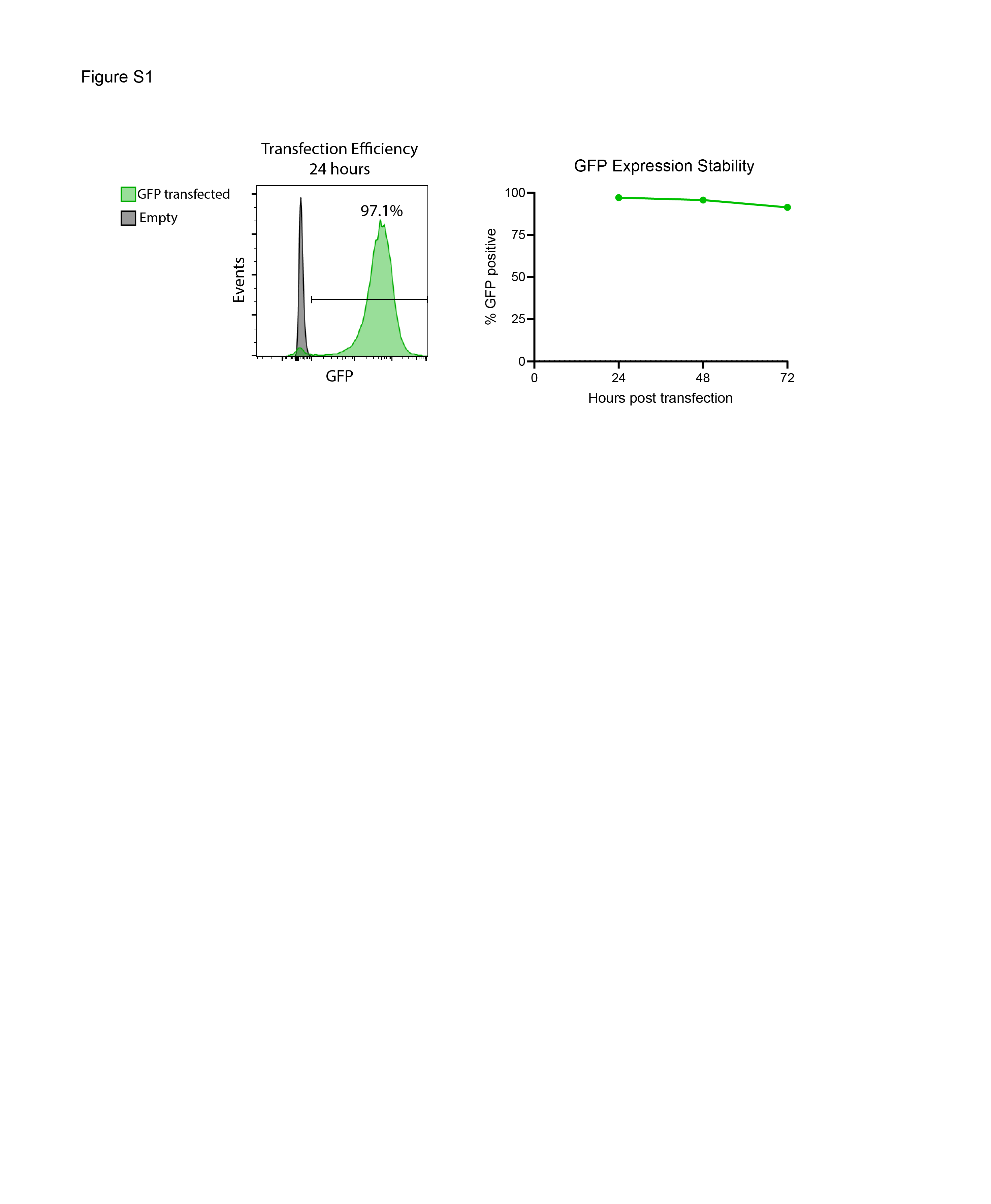

### Supplemental Figure 2

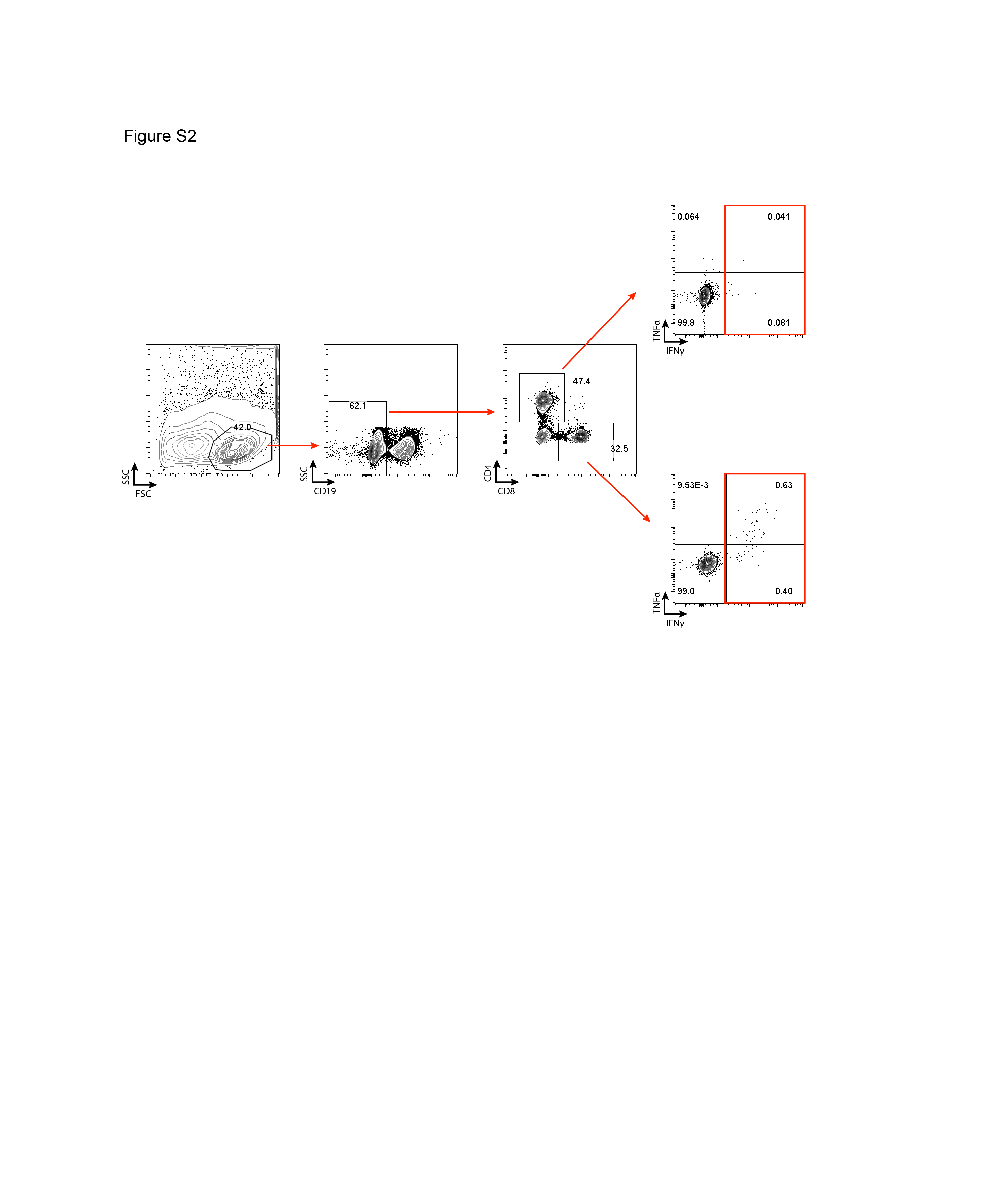

### Supplemental Figure 3

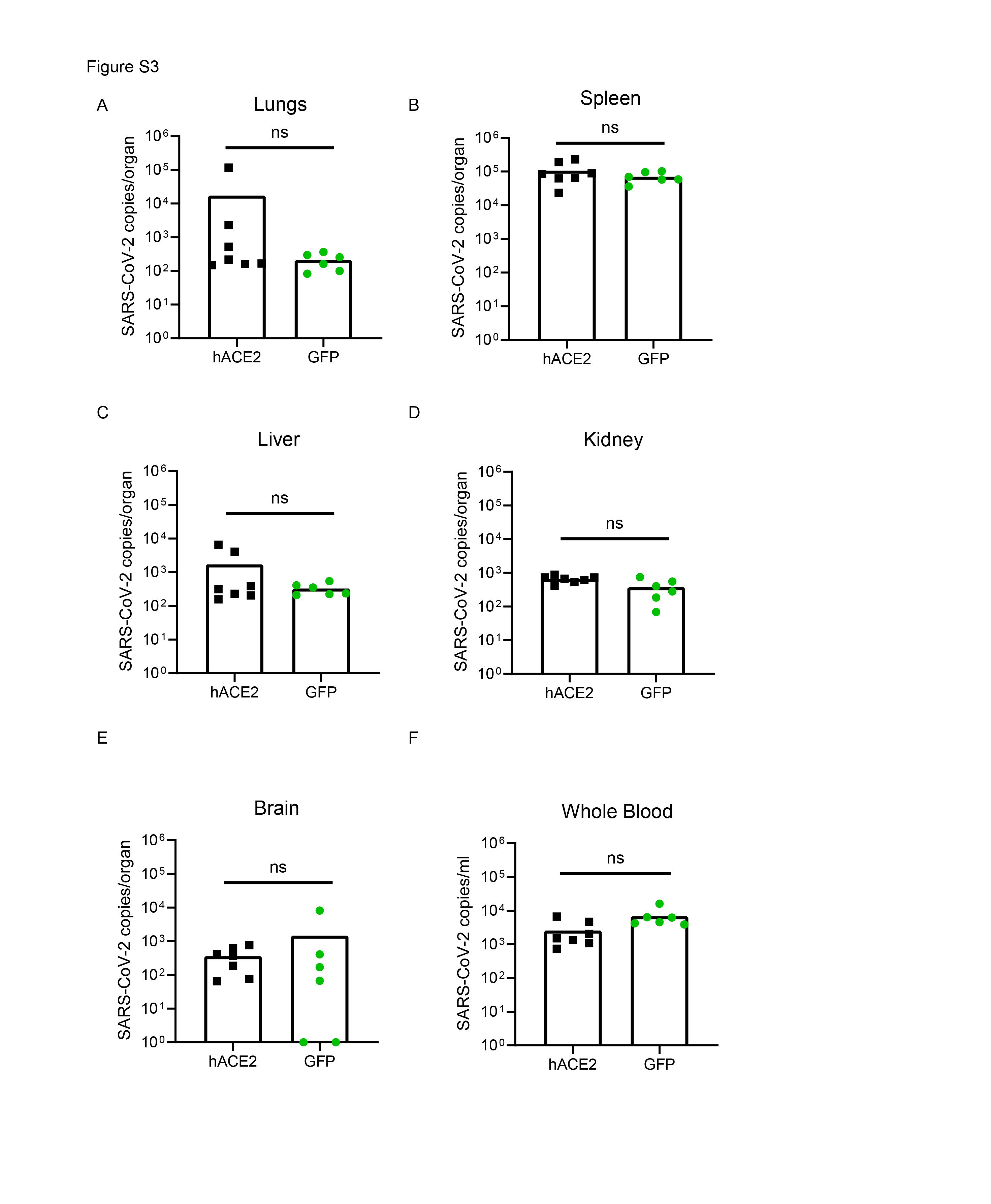

### Supplemental Figure 4

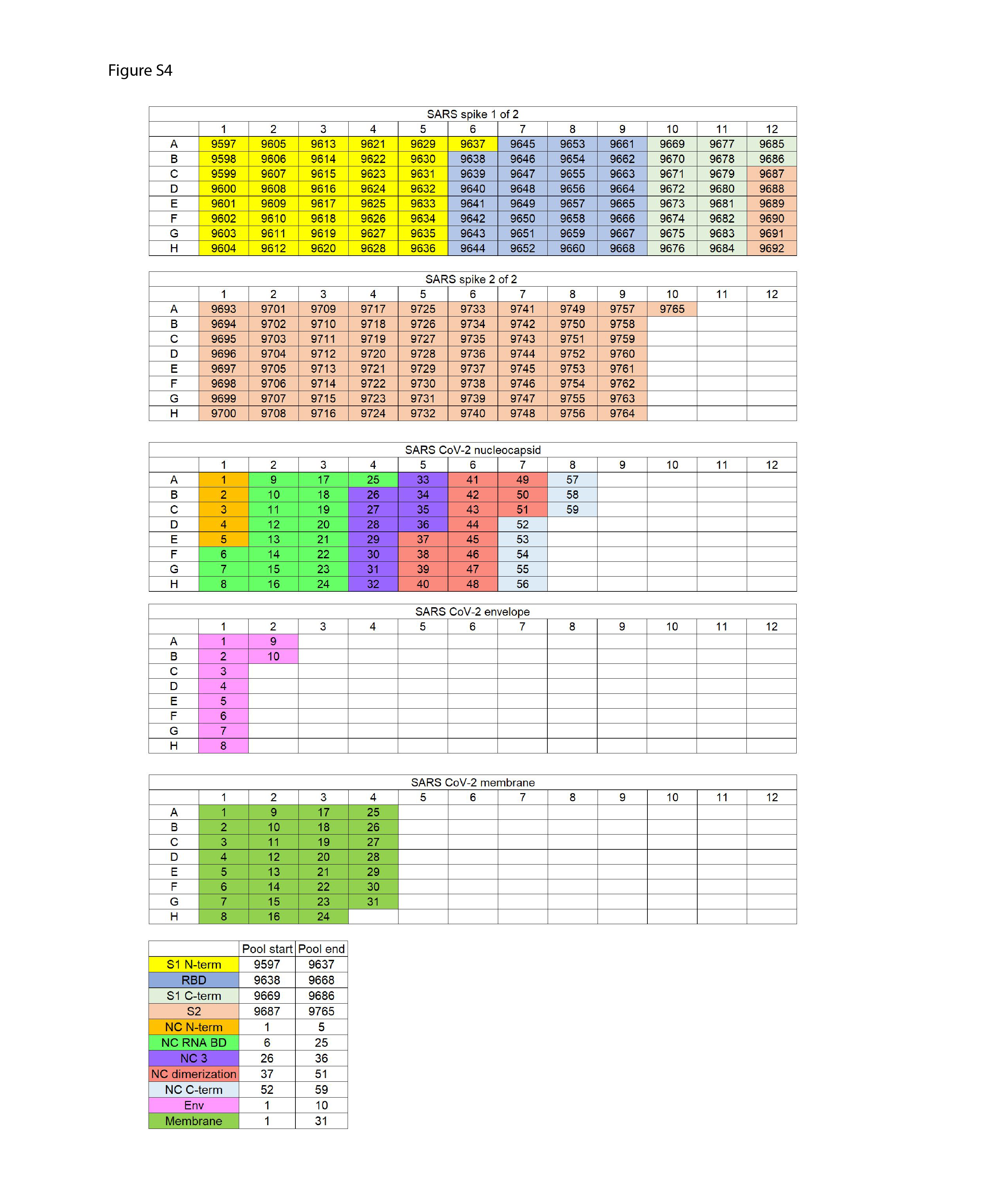
